# ToxINsight: An Integrated Web Platform for Sequence, Structure, and Ligand-Binding Analysis of Snake Venom Toxins

**DOI:** 10.64898/2026.10.27.762231

**Authors:** Aayushi Gupta, Tukkapuram Venkata Lalith Krishna, Bipin Singh

## Abstract

Around 5.4 million people worldwide are bitten by snakes each year, with 1.8 to 2.7 million cases of envenomation, of which 81,410 to 137,880 people die from snake envenomation worldwide each year. It is categorized as a high-priority Category A Neglected Tropical Disease (NTD) by the World Health Organization (WHO). A molecular level understanding of snake venom toxins and their mechanisms of action is crucial for development of efficient therapeutics tackling snake envenomation. Snake venoms are complex mixtures of bioactive peptides and proteins that have also served as templates for drug discovery. Resources dedicated to snake venom toxins sequence, structure, functions, geographical distributions and molecular-level analysis remain fragmented across general-purpose repositories. Here we present ToxINsight, an integrated, freely accessible database and web-based analysis platform built exclusively for snake venom toxins. ToxINsight curates 2,078 toxin entries spanning 11 medically and scientifically relevant snake toxin families across 237 venomous snake species from five different snake families, and provides 2,540 three-dimensional structures associated with these entries, which are assembled from the RCSB Protein Data Bank, AlphaFold DB and ESMFold-generated 3D models. ToxINsight offers built-in tools alongside toxin records for comparisons and analysis of toxin sequences and structures and their properties for efficient target selection and prioritization. ToxINsight allow the user to select multiple entries and align them, or select a pair of entries to compare structural and physicochemical properties. ToxINsight provides all reported-ligand annotations for each toxin entry, and bulk export of sequence, structure, and metadata for building sequence and/or structure based machine learning models. ToxINsight also offers a visual ATLAS, which provides geographical information about venomous snake families/species for a more focused understanding of snake venom toxins across different geographical landscapes. By combining sequence, structure, ligand and geographical information along with analysis and comparative tools in a single interface, ToxINsight is designed to support and encourage researchers across the world towards the development of efficient targeted therapeutics against snake venom toxins from diverse families. It provides an interface for integrated understanding toxin sequences, structures and physicochemical properties, for providing scaffolds for designing antivenom and for comparative venomics. ToxINsight is freely accessible at https://toxinsight.in/.

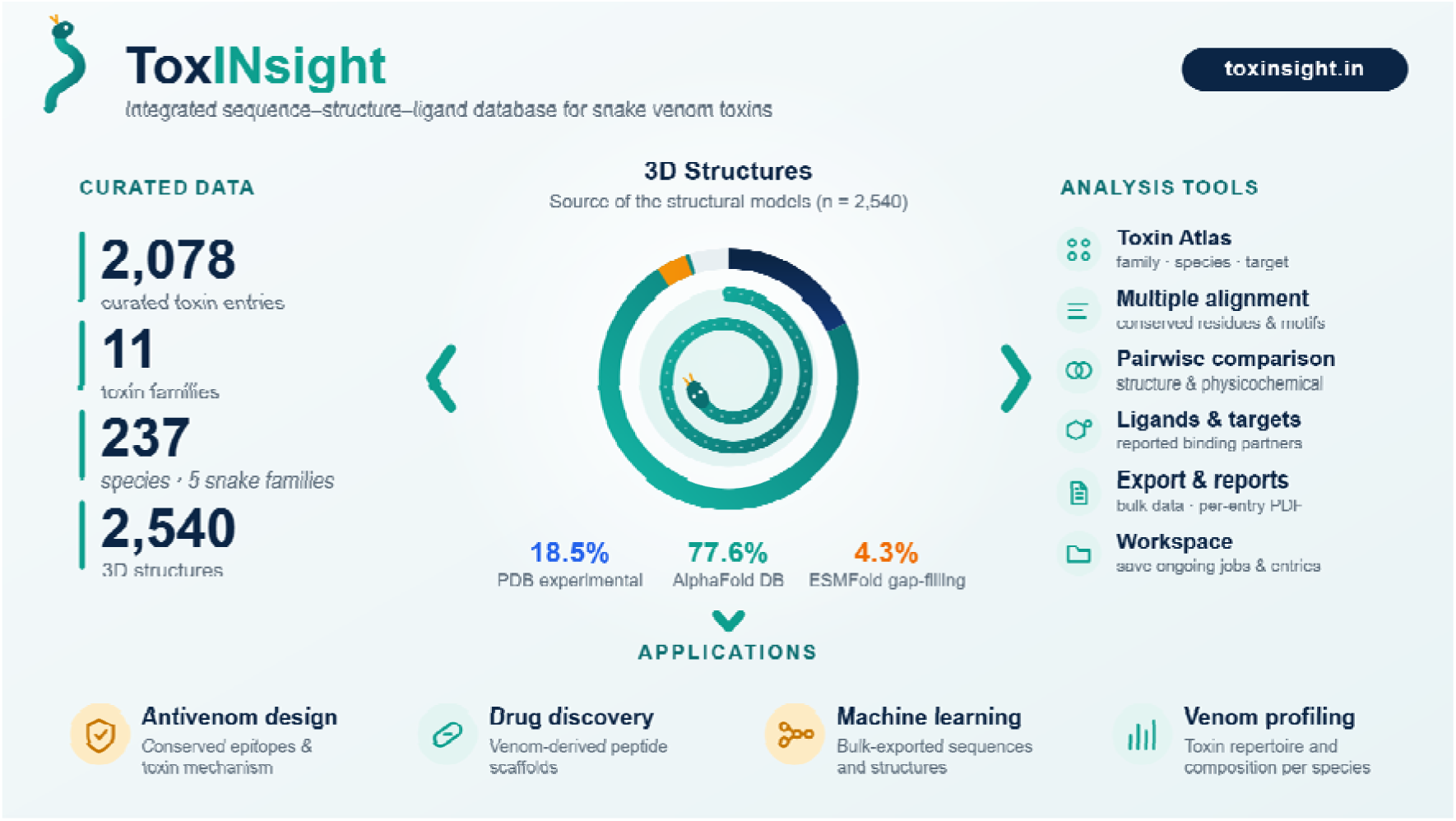

## 1. INTRODUCTION

Snake envenomation has been classified under tropical neglected disease since 2017 by WHO. Millions of lives are affected by snake envenomation, out of which 81,410 to 137,880 people face fatal consequences, and around 400,000 people survive envenomation each year but are left with permanent disabilities or amputations^1^. Molecular level exploration of snake venom toxins and their underlying mechanisms of action is key to designing effective drugs to counter envenomation and is a high priority. Currently, the available antivenoms are animal-derived polyclonal antibodies that are constantly losing their efficiency due to evolutionary changes in venom variation and composition over time, and they also result in severe side effects due to their use in high doses. These antivenoms are not able to cope with the constantly evolving nature of snake venom toxins found in native venomous snakes, as these antivenoms are prepared from captive snakes^2–4^.

Snake venoms are constantly evolving and contain a complex mixture of peptides and proteins^5^. Some of the proteins in snake venoms have evolved to have remarkable potency and target specificity for either ion channels, receptors (like nAChRs) or coagulation-cascade enzymes. This combination of potency and selectivity has made snake venom toxins a deadly threat and also an enduring source of pharmacological tools and clinically approved drugs. Snake venom-derived drugs that are clinically approved are in use today (Table 1), and additional venom-derived peptides continue to move through preclinical and clinical development.

**Table 1.**
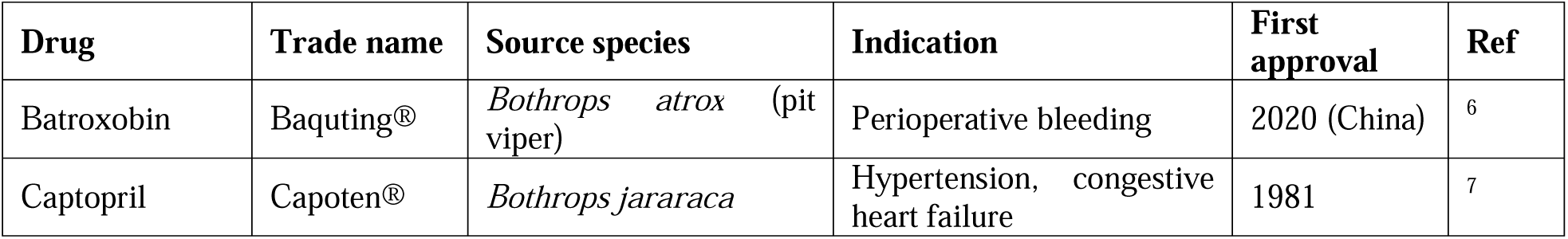
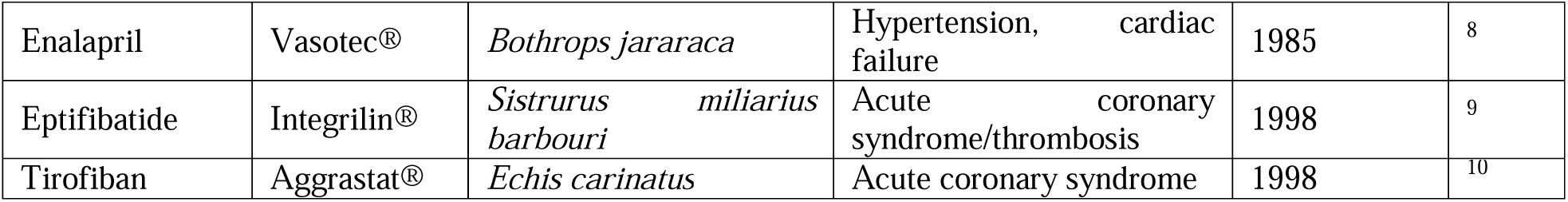
Selected snake venom-derived drugs approved for clinical use.

| **Drug** | **Trade name** | **Source species** | **Indication** | **First approval** | **Ref** |
|---|---|---|---|---|---|
| Batroxobin | Baquting® | *Bothrops atrox* (pit viper) | Perioperative bleeding | 2020 (China) | <sup>6</sup> |
| Captopril | Capoten® | *Bothrops jararaca* | Hypertension, congestive heart failure | 1981 | <sup>7</sup> |
| Enalapril | Vasotec® | *Bothrops jararaca* | Hypertension, cardiac failure | 1985 | <sup>8</sup> |
| Eptifibatide | Integrilin® | *Sistrurus miliarius barbouri* | Acute coronary syndrome/thrombosis | 1998 | <sup>9</sup> |
| Tirofiban | Aggrastat® | *Echis carinatus* | Acute coronary syndrome | 1998 | <sup>10</sup> |

Designing and optimizing venom-derived therapeutics, and in-depth molecular understanding of venom toxins, will require access not only to toxin sequences but their three-dimensional structures, physicochemical properties and their reported bound ligands. General-purpose protein and ligand/drug resources such as UniProt^11^, the RCSB Protein Data Bank (PDB)^12,13^, ChEMBL^14,15^, and DrugBank^16^ contain toxin-relevant records as part of much broader collections, while dedicated venom-toxin resources such as ConoServer^17^ (cone snail conopeptides), ArachnoServer^18^ (spider toxins), and Kalium^19^ (potassium-channel-acting peptides) focus on other venomous lineages or on a single target class. Among snake venom-focused resources, VenomZone^20^ (from Expasy) curates functional annotation for a broad range of venom protein families, which includes venom protein families from snakes, and a general cross-taxon biotoxin database^21^. However, none of these resources is built specifically around snake venom toxins and lacks integrated structural information, relevant physicochemical properties and geographical coverage. Further, these resources do not provide an integrated set of tools for analyzing sequence similarity, multi-toxin alignment, and pairwise structural/physicochemical comparison under a single interface.

To address this gap and bring all the relevant snake venom toxin data together in one place, we designed ToxINsight, a snake venom toxin database that combines curated sequences, structural features, ligand and geographical coverage with a set of analysis and comparative tools across 6 different medically relevant venomous snake families. ToxINsight currently contains 2,078 toxin entries across 11 toxin families, drawn from 237 species representing five snake families, together with 2,540 associated three-dimensional structures. In this article, we describe data curation of snake venom toxin sequences, structures, and associated ligands, along with the pipeline behind ToxINsight. We also describe key applications of ToxINsight and the analysis tools available through its web interface (Atlas, multiple sequence alignment (MSA), pairwise structural and physicochemical properties comparison, and bulk export), and a set of representative practical use cases illustrating how ToxINsight will support sequence-structure–activity relationship analysis of snake venom toxins, comparative venomics, venom-derived drug discovery and machine learning based predictive model development

## 2. MATERIALS AND METHODS

### 2.1. Data Collection and Curation

We collected ToxINsight data from open-source databases. This includes snake venom toxin information from eleven toxin families that contribute significantly to the toxicity of snake venoms. The sequence information was obtained from the snake venom toxin families documented in Venomzone ^20^, which links the toxin families to their respective UniProt ^11^ entries (Swiss-Prot and TrEMBL). Protein sequences, functional annotations, taxonomy information, GO terms ^22,23^, and references to the literature were all retrieved from UniProt^11,24^. Experimentally determined structures were taken from the RCSB Protein Data Bank ^12^, while predicted structures were from the AlphaFold Protein Structure Database^25,26^, together with their per-residue pLDDT values. ESMFold^27^ was used to predict and document structures for entries without them. We have collected interaction data for each entry available from UniProt and IntAct^28^, which are interconnected resources that collectively map the landscape of protein sequences, functional associations, and physical molecular interactions. The binding pockets were calculated using the DoGSiteScorer REST API^29,30^. Ligand information for the entries was taken from BindingDB^31^ (based on measured binding affinities) and co-crystallized ligands with the PDB structures.

### 2.2. Online Implementation and Analysis Toolset

ToxINsight is a full-stack web application whose frontend was created with React 19^32^, TypeScript^33^, Vite^34^, and Tailwind CSS v4^35^. Structures of the proteins can be visualised in the browser with PDBe Molstar plugin^36^. The backend was built with Python^37^ and FastAPI^38^, whereas PostgreSQL 16^39^ was used as our database management system. Database interactions were handled with SQLAlchemy 2.0^40^, while Alembic^41^ was used for database schema migrations. Thus, the structural data is available for more than 98% of proteins in our dataset. Physicochemical properties of the proteins were calculated based on their sequences by the ProtParam^42^ module in Biopython.

The prediction of binding pockets was achieved using the DoGSite3^43^ service available at ProteinsPlus^30^, and structural motifs were detected using Foldseek^44^ and Folddisco^45^. Information regarding protein–protein interactions was gathered from UniProt and IntAct, while ligand information was extracted from BindingDB and ChEMBL. Additionally, the platform features a geographic visualization of the dataset, including the distribution of 237 venomous species in 124 countries, via an interactive D3.js globe^46^.

The relational schema of ToxINsight is shown in Figure 1. The protein table is the central entity and contains the UniProt accession, name, sequence, length, review status and function for each toxin entry. All other information is linked to the central entity through separate tables. Each protein belongs to one species and one toxin family, which have their own tables so that taxonomic browsing and toxin family-based filters work smoothly. A protein/toxin entry can have more than one structure, such as experimental entries and predicted models, so structures form a one-to-many relation. Each structure record keeps its own source, method, resolution and pLDDT score. This is why the 2,540 structures outnumber the 2,078 total toxin entries. Physicochemical properties are stored once per protein, whereas reported ligands, interaction partners and Gene Ontology terms can each be multiple, so they are one-to-many relations. Predicted binding pockets and structural similarity results are stored internally as JSON in their own tables, each with a status and job identifier, and are displayed to users as formatted tables in the interface. References are linked through a junction table, since one publication can describe several toxins and one toxin can be described in several publications. This structure allows entry pages, workspace collections and bulk exports to be assembled from the same underlying records.

**Figure 1.**
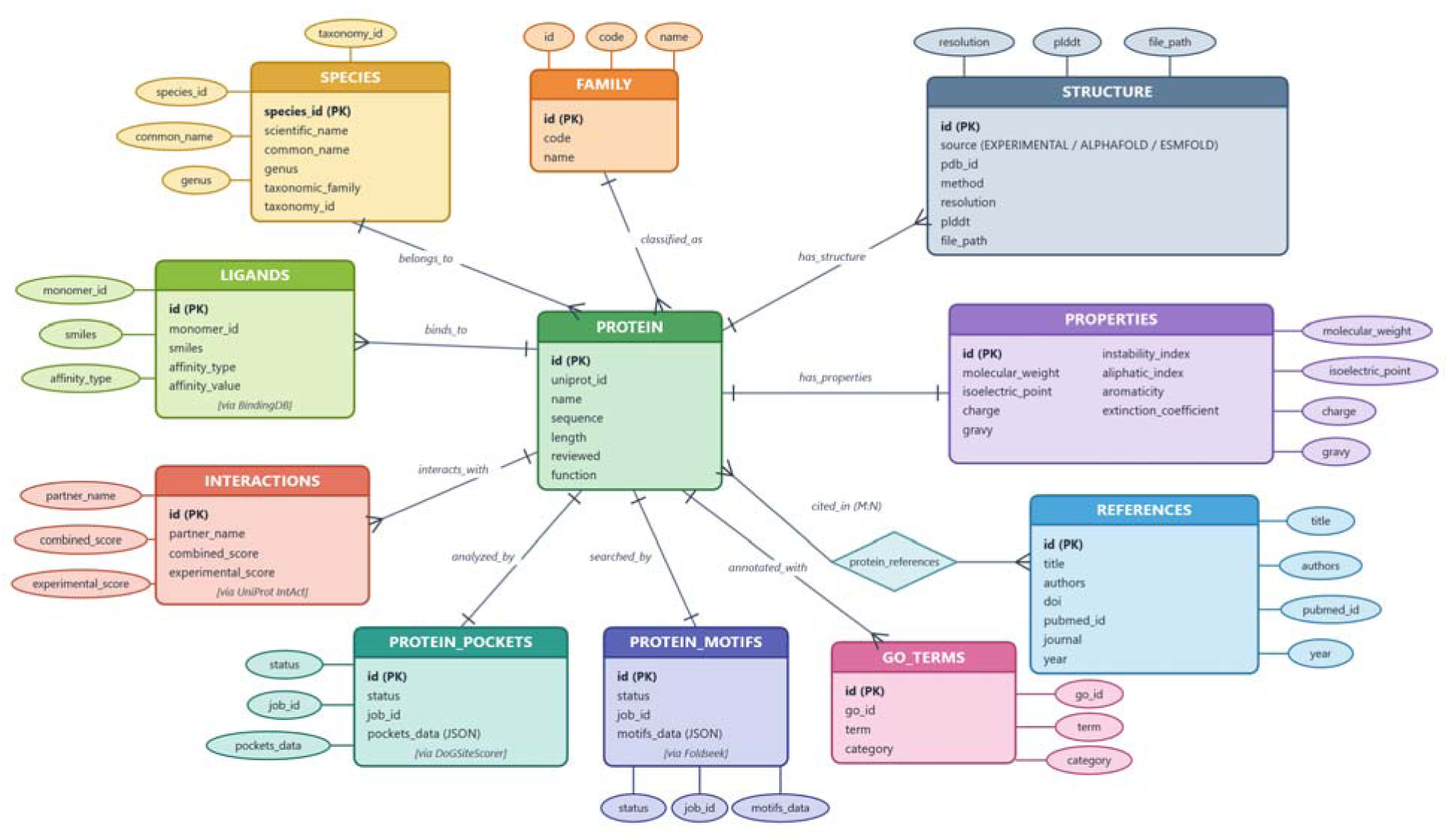
Relational Database Architecture. The schema organizes 11 tables around the primary PROTEIN entity: SPECIES (N:1), FAMILY (N:1), STRUCTURE (1:N, PDB & AlphaFold), PROPERTIES (1:1, MW, pI, GRAVY, instability), GO_TERMS (1:N), REFERENCES (M: N join), PROTEIN_POCKETS (1:1, DoGSiteScorer predictions), PROTEIN_MOTIFS (1:1, Foldseek structural alignments), INTERACTIONS (1:N, UniProt IntAct interactome), and LIGANDS (1:N, BindingDB small-molecule binders).

## 3. RESULTS AND DISCUSSION

### 3.1 ToxINsight metrics

ToxINsight has a robust and complete collection of all the snake venom toxins which are medically relevant for snake venom toxicity (Figure 2B). The data have been classified and processed based on venomous snake families, venom toxin families and structural properties (Figure 2). A total of 2078 toxin entries (Table 2) with 2540 structures were collected across 237 venomous snake species. These data include five venomous snake families from the suborder Serpentes. From a total of 237 species, 128 were Viperidae, 88 were Elapidae, 12 were Colubridae, 2 were Lamprophiidae, 2 were Homalopsidae, and 5 were unclassified (Figure 2C). From all 2078 toxin entries taken from UniProt, 1932 (93.0%) were from Swiss-Prot (reviewed), and 146 (7.0%) were from TrEMBL (unreviewed) (Figure 2D). Only entries with a defined amino acid sequence and a traceable literature or database source were retained. For each entry, we recorded the toxin name and synonyms, source species and family, UniProt accession ID, gene annotation records, full-length and signal-peptide-cleaved sequence, molecular mass, and, if reported, the disulfide-bond connectivity. We have selected 11 prominent toxin families in our database that are known to be medically relevant for snake venom toxin research purposes (Figure 2A). From these 2078 toxin entries, 537 were 3FTX, 508 were PLA_2_, 272 were SVMP, 220 were CTL, 219 were SVSP, 152 were KUN, 149 were M15, 7 were DIS, 5 were LAAO, 5 were CRISP, and 4 were NP. (Figure 2A and Table 2)

**Figure 2.**
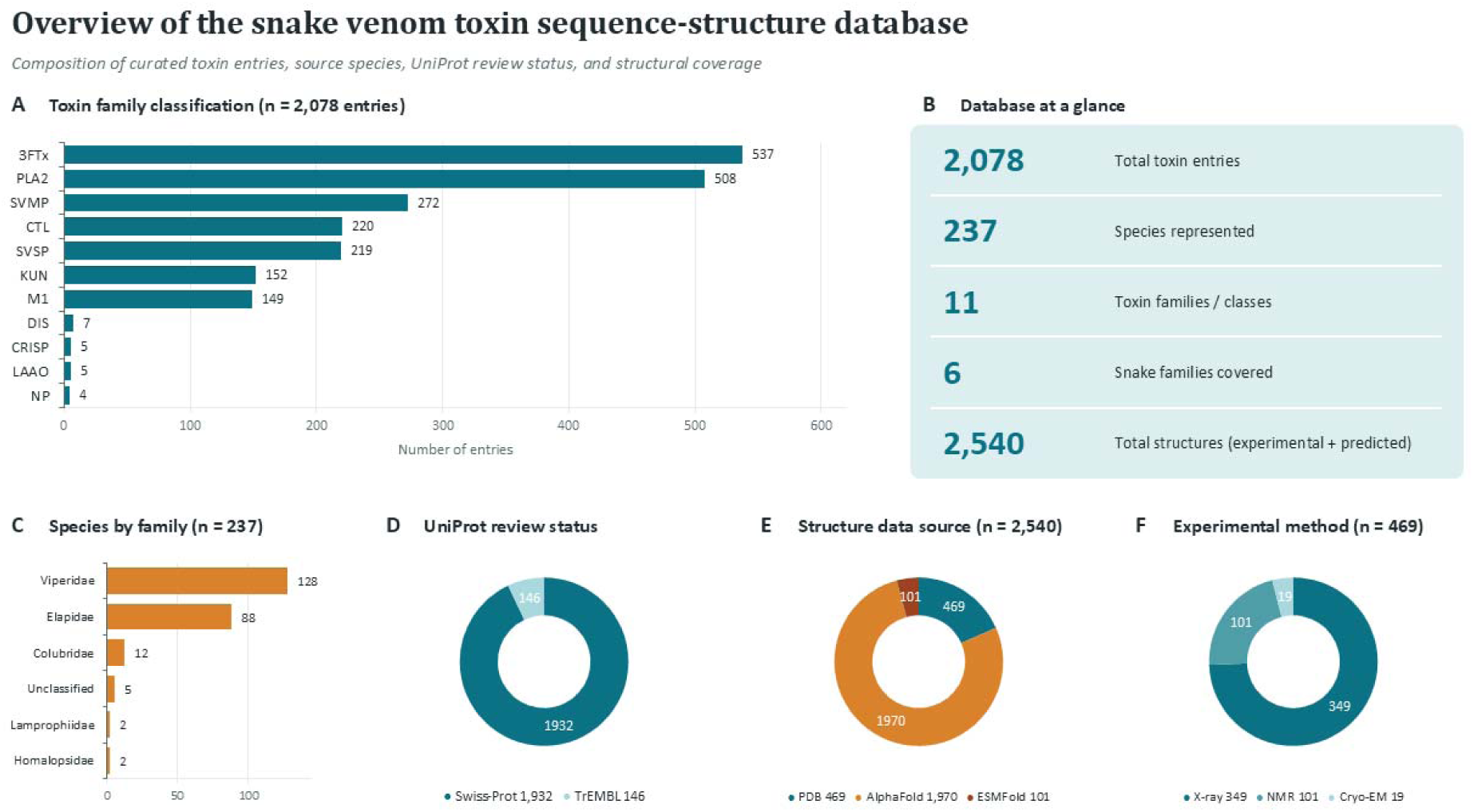
Overview of ToxINsight content. (A) Toxin family classification (n = 2,078 entries). (B) Database-at-a-glance summary statistics. (C) Species distribution by snake family (n = 237 species). (D) UniProt review status of curated entries. (E) Source of the three-dimensional structure associated with each entry (n = 2,540 structures). (F) Experimental method breakdown for the 469 PDB-derived structures in (E).

**Table 2.** Toxin family classification used in ToxINsight.

| **Abbreviation** | **Toxin family** | **Entries (n)** |
|---|---|---|
| 3FTx | Three-finger toxin | 537 |
| PLA2 | Phospholipase A2 | 508 |
| SVMP | Snake venom metalloproteinase | 272 |
| CTL | C-type lectin / lectin-like protein | 220 |
| SVSP | Snake venom serine protease | 219 |
| KUN | Kunitz-type protease inhibitor | 152 |
| M15 | Peptidase M1 | 149 |
| DIS | Disintegrin | 7 |
| CRISP | Cysteine-rich secretory protein | 5 |
| LAAO | L-amino-acid oxidase | 5 |
| NP | Natriuretic peptide | 4 |

From the total of 2540 structures available, 469 were experimentally resolved structures, and 1970 were predicted from AlphaFoldDB (Figure 2E). Out of 469 PDB entries, 349 were obtained using X-ray crystallography, 101 were obtained using NMR, and 19 were obtained using Cryo-EM (Figure 2F). Thus, the structural data are available for more than 98% of proteins in our dataset. Apart from these structures, the missing structural information for 101 data records, which mainly pertains to fragments, was predicted using ESMFold.

### 3.2. Distribution ATLAS

ToxINsight provides a visual Atlas for the user to browse and visualize countries across the world for venomous snake families and species. Users can visualize venomous snake species/family richness of a country or region (Figure 3A), select a specific venomous snake family to highlight all the countries it is reported from (Figure 3B), and users can also select or search a specific venomous snake species name or their common name to highlight all the countries it has been reported from (Figure 3C). Atlas also provides a flat and globe view to represent the world map as per the user’s preference (Figure 3D). This browsing layer is intended to support exploratory analysis, hypothesis-generation (e.g., “show me every 3FTx reported to act on nicotinic acetylcholine receptors”) as well as targeted lookup of a specific and previously identified snake venom toxin.

**Figure 3.**
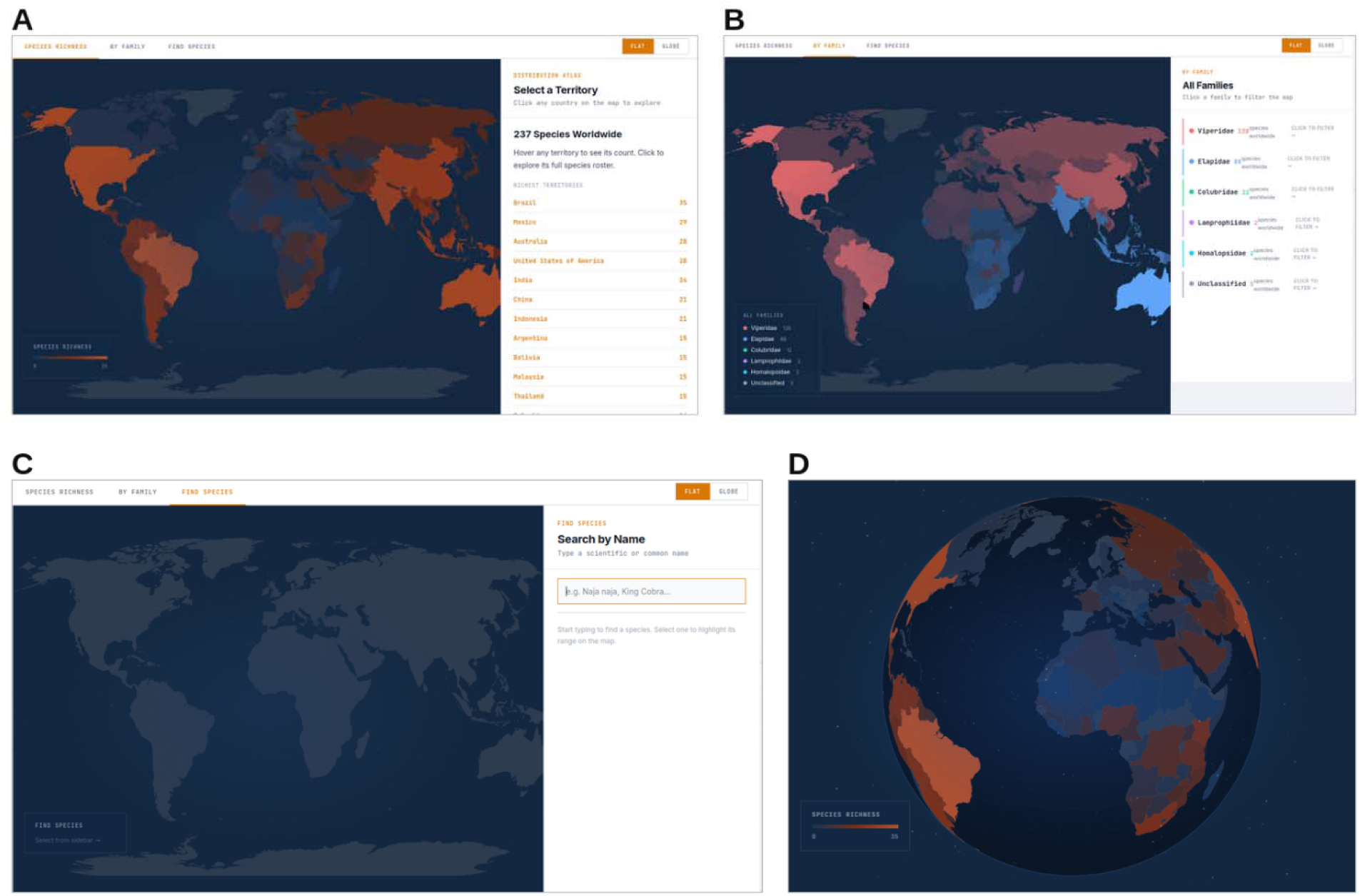
Distribution Atlas: (A) Species richness per country. (B) Venomous snake family-based view. (C) Species search. (D) Globe view.

### 3.3. Browsing ToxINsight

ToxINsight allows the user to search the database based on taxonomic inputs using their venomous snake families, genera, and species (Figure 4B), which can be found in the taxonomy section provided in the sidebar navigation section (Figure 4A). ToxINsight offers a keyword-based search across the platform, where users can search by toxin name, synonyms, UniProt identifiers, species name, common name, and reported targets, using fuzzy words and fuzzy-string matching to retrieve relevant results. Apart from searching using terms in the search bar, users can navigate to the search page under the dashboard for a more focused search using filters to organise their query results according to toxin families, taxonomic family, species, range for toxin length (Figure 4D), and the user can also set the requirement for the toxin entries in the results to have a 3D structure or to be reviewed (Swiss-Prot) (Figure 4C). User have the option to save toxin entry selections to the workspace to save their work and progress (Figure 4E). These saved entries can then be found in the workspace section of the index. After searching through the database as per the user’s selections, the user can then go through each entry for their respective toxin information. The user can download individual toxin information in PDF form or can select multiple entries to download their FASTA sequences in bulk. User-selected sets of toxins can be exported in bulk as sequence (FASTA), structure (PDB), or tabular (CSV) files, supporting downstream use in predictive ML model building, local bioinformatics pipelines, docking or virtual screening workflows, and phylogenetic analysis.

**Figure 4.**
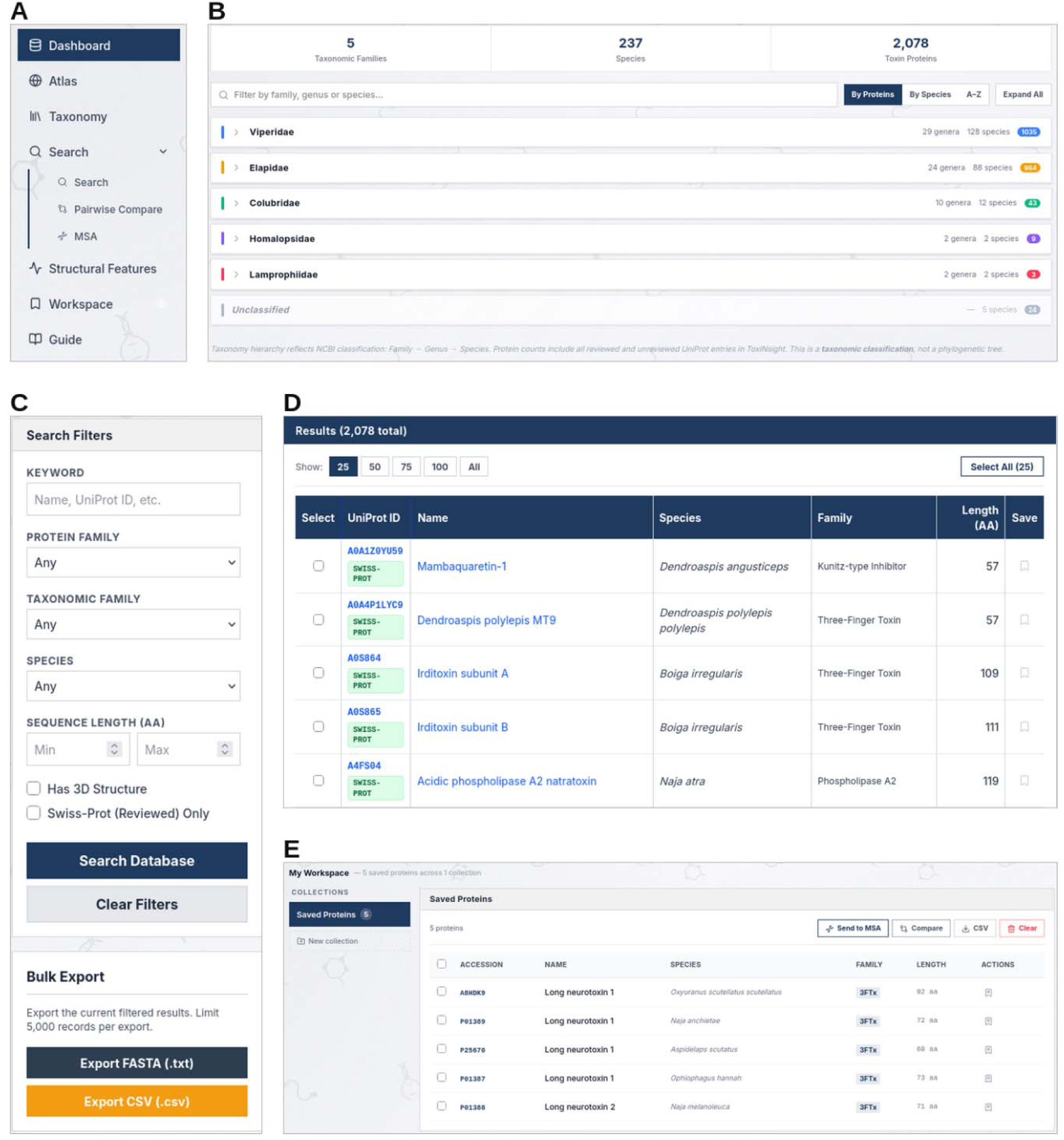
Browsing and searching ToxINsight. (A) Taxonomy browser. (B) Sidebar navigation/ index. (C) Search filters and bulk export option. (D) Results table view. (E) Workspace of saved entries.

### 3.4. Toxin Entry Pages: Sequence, Structure, and Properties

Selecting a specific toxin entry will display toxin information to the user. This toxin information comprises the toxin name, toxin family, amino acid length (AA), source organism, UniProt ID, UniProt Status (Reviewed/Unreviewed), number of 3D structures available with their source or related references (for experimental structures), 3D structure view of these structures, along with ESMFold for structure prediction, physicochemical properties, amino acid sequence, Gene Ontology data, and literature references linked with the corresponding entry. The physicochemical properties include molecular weight (Da), Isoelectric point (pI), GRAVY index, instability index, aliphatic index, aromaticity, and Extinction coefficient (Figure 5A). The amino acid sequence display section highlights amino acids according to their charge, polarity and hydrophobicity. Gene annotation information is provided under the Gene Ontology (GO) section (Figure 5B). At the top of the page, users get the option to download the FASTA sequence, structure, toxin report and save the entry to the workplace section for later access (Figure 5B). At the bottom of the toxin information, users also get the link to the entry’s structural feature information, which, when prompted, will display calculated binding pocket details, predictions on structural similarity metrics and interactions, along with any reported ligands for the toxin entry. This design mirrors, for a single snake-focused toxin universe, the kind of consolidated sequence–structure–activity record that general biotoxin resources provide across much broader and more heterogeneous taxonomic scope.

**Figure 5.**
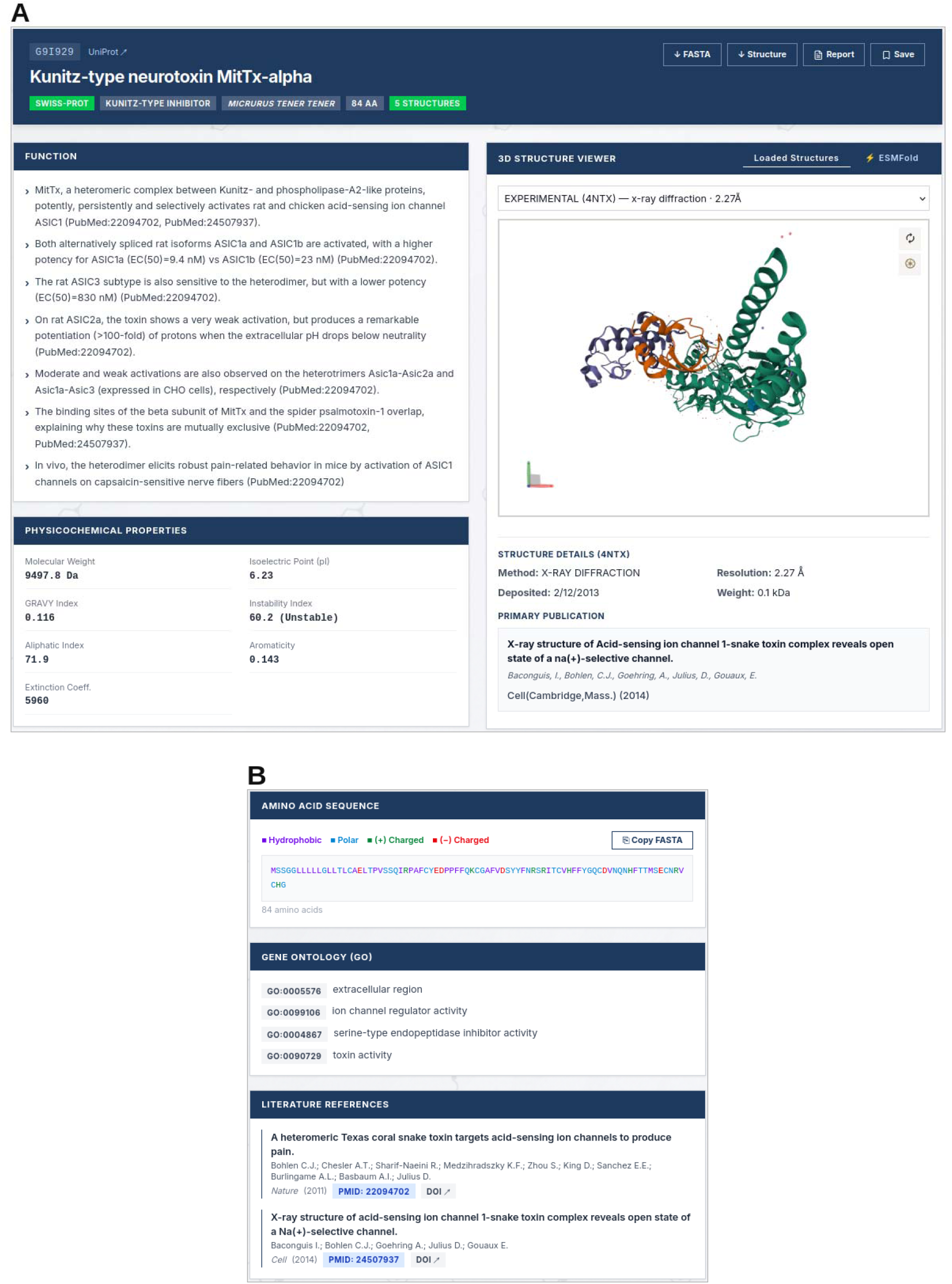
Toxin entry page (example: G9I929, Kunitz-type neurotoxin MitTx-alpha, Micrurus tener tener). (A) Entry header with download and save options, functional description, physicochemical properties, and the 3D structure viewer with experimental structures (PDB 4NTX shown) and an ESMFold tab. (B) Colour-coded amino acid sequence, Gene Ontology annotations and linked literature references.

### 3.5 Structural features

The structural features page includes four sections: binding pocket, structural similarity, interactions, and reported ligands, with their availability displayed for toxin entries together (Figure 6A). The binding pocket section comprises features for binding pocket/s predicted using DoGSiteScorer, which includes binding pocket volume (Å^3^), surface area (Å^2^), drug score (composite descriptor) and druggability (ML-based prediction of how likely the pocket is to bind with a drug molecule) (Figure 6B). The structural similarity predicted by FoldSeek provides the top 15 structural hits, which can be selected to display their structure along with their database, probability (statistical confidence that the reported alignment is not random), E-value (the expected number of alignments with a given score that would occur by chance), sequence identity (fraction of aligned residues that are identical between query and target) and aligned length (the span of the matched region). The interaction data gathered from IntAct and UniProt include partner gene/s, UniProt ID, source (UniProtKB/IntAct/BioGRID/STRING/DIP), score (the strength or quality of the interaction) and confidence (indicating how confident the database is that the interaction is real).

**Figure 6.**
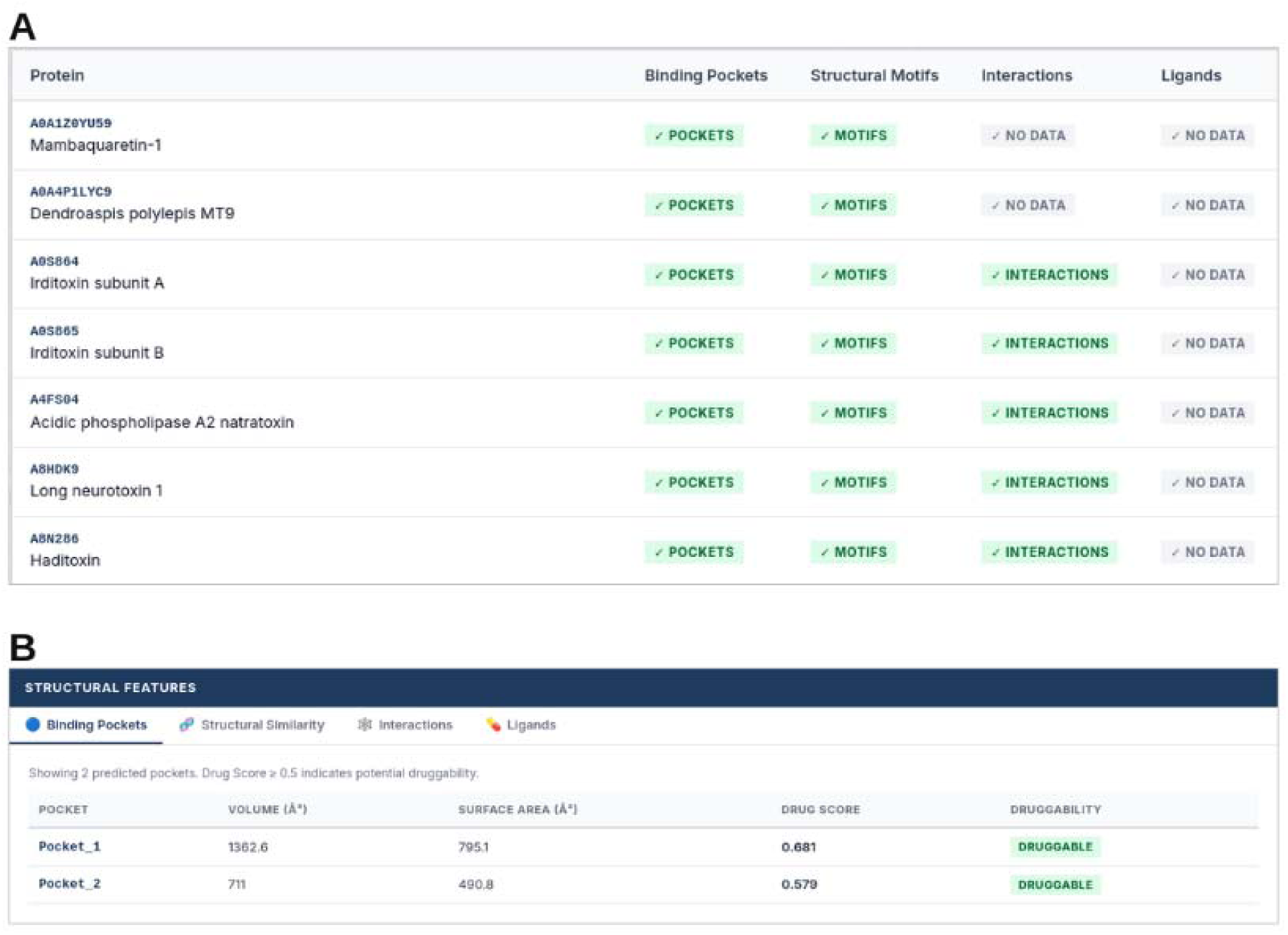
Structural features. (A) Availability of binding pockets, motifs, interactions and ligands per entry. (B) Predicted binding pockets with volume, surface area, drug score and druggability; structure similarity, interactions and ligands.

### 3.6. Multiple Sequence Alignment

ToxINsight supports selection of multiple toxin entries for multiple sequence alignment, allowing users to visualize alignments of the selected sequences, conserved residues, motifs, and lineage-specific substitutions (Figure 7). MSA is performed using Clustal Omega^47^, which is based on HMMs^48^ (Hidden Markov Models) and mBed^49^ clustering.

**Figure 7.**
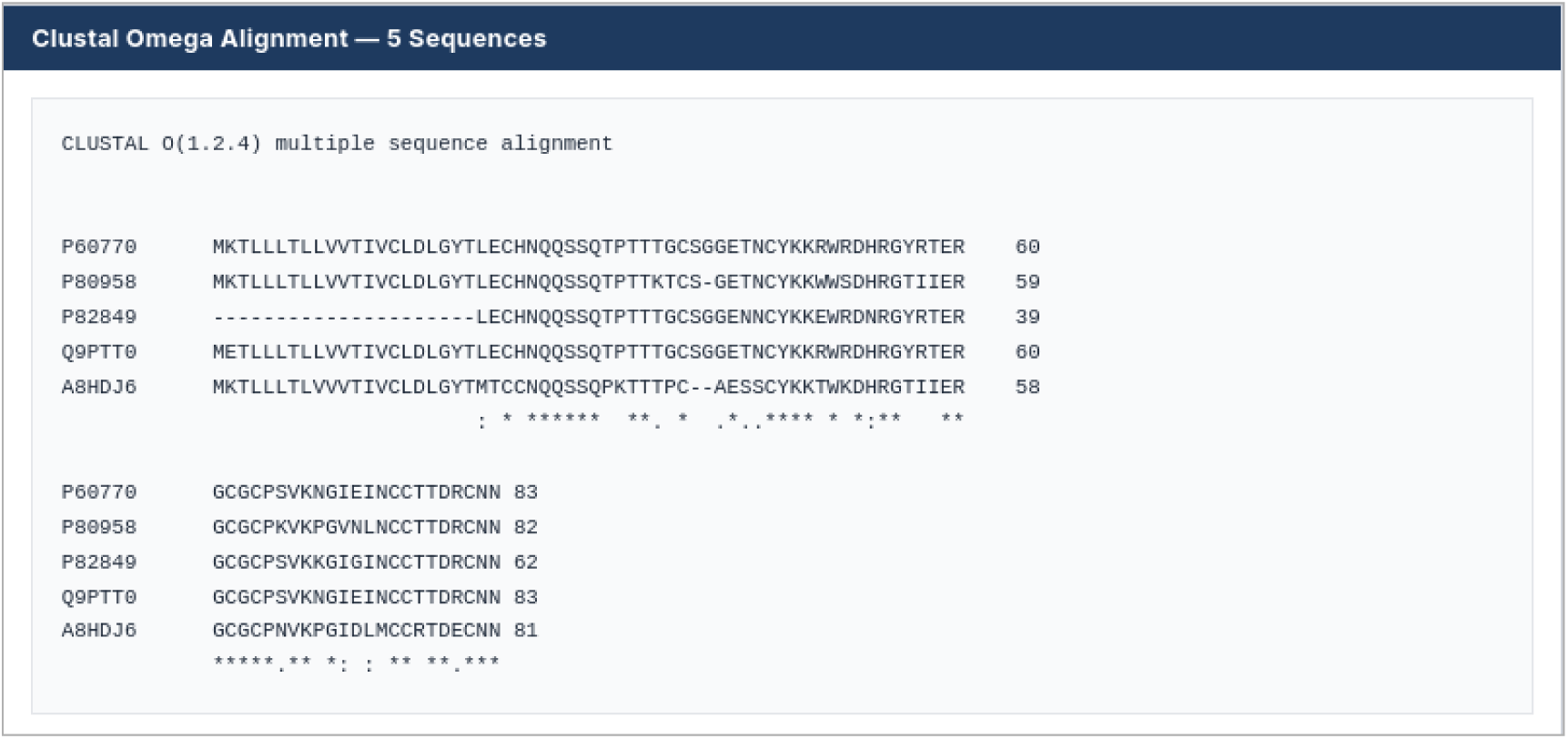
Multiple sequence alignment. Clustal Omega alignment of five selected toxins as an example.

### 3.7. Pairwise Structural and Physicochemical Comparison

ToxINsight offers pairwise comparisons of two selected toxin entries, primarily based on their structure and physicochemical properties. The comparison is made between selected toxin entries’ information, such as species, toxin family, toxin length, mass (Da) and pI (Figure 8A), followed by comparing physicochemical properties with a visual representation (Figure 8B) and 3D structure comparison (superimposed or side-by-side) (Figure 9A and 9B). In addition, it globally aligns both toxin sequences and reports the alignment and sequence identity between the two (Figure 8C). This comparison is designed to help distinguish cases where change in sequence between two toxins is reflected in their structural or physicochemical properties from cases where the overall fold and properties are conserved despite their sequence differences.

**Figure 8.**
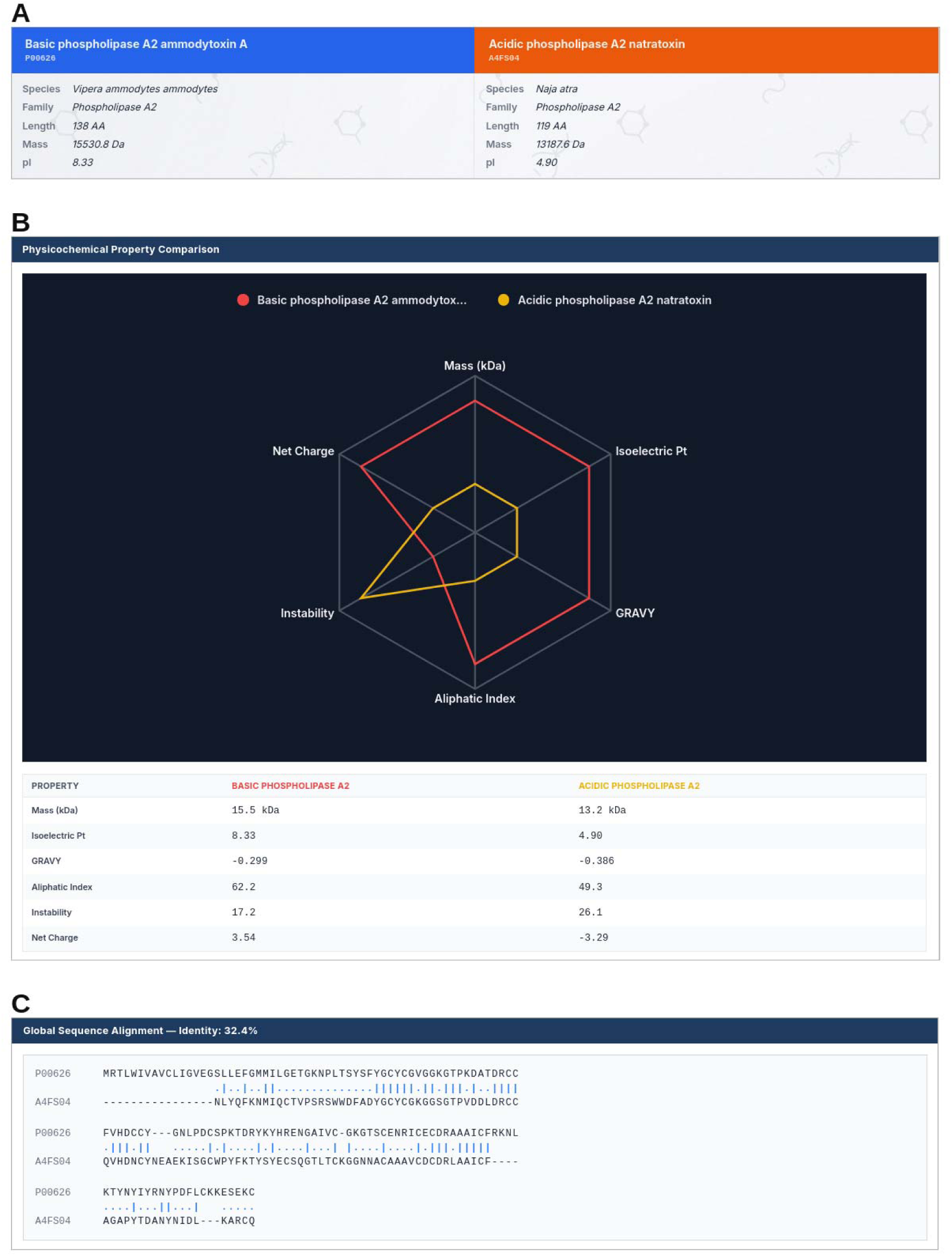
Pairwise comparison (example: P00626 and A4FS04). (A) Entry summary. (B) Physicochemical property radar chart and table. (C) Global sequence alignment (32.4% identity).

**Figure 9.**
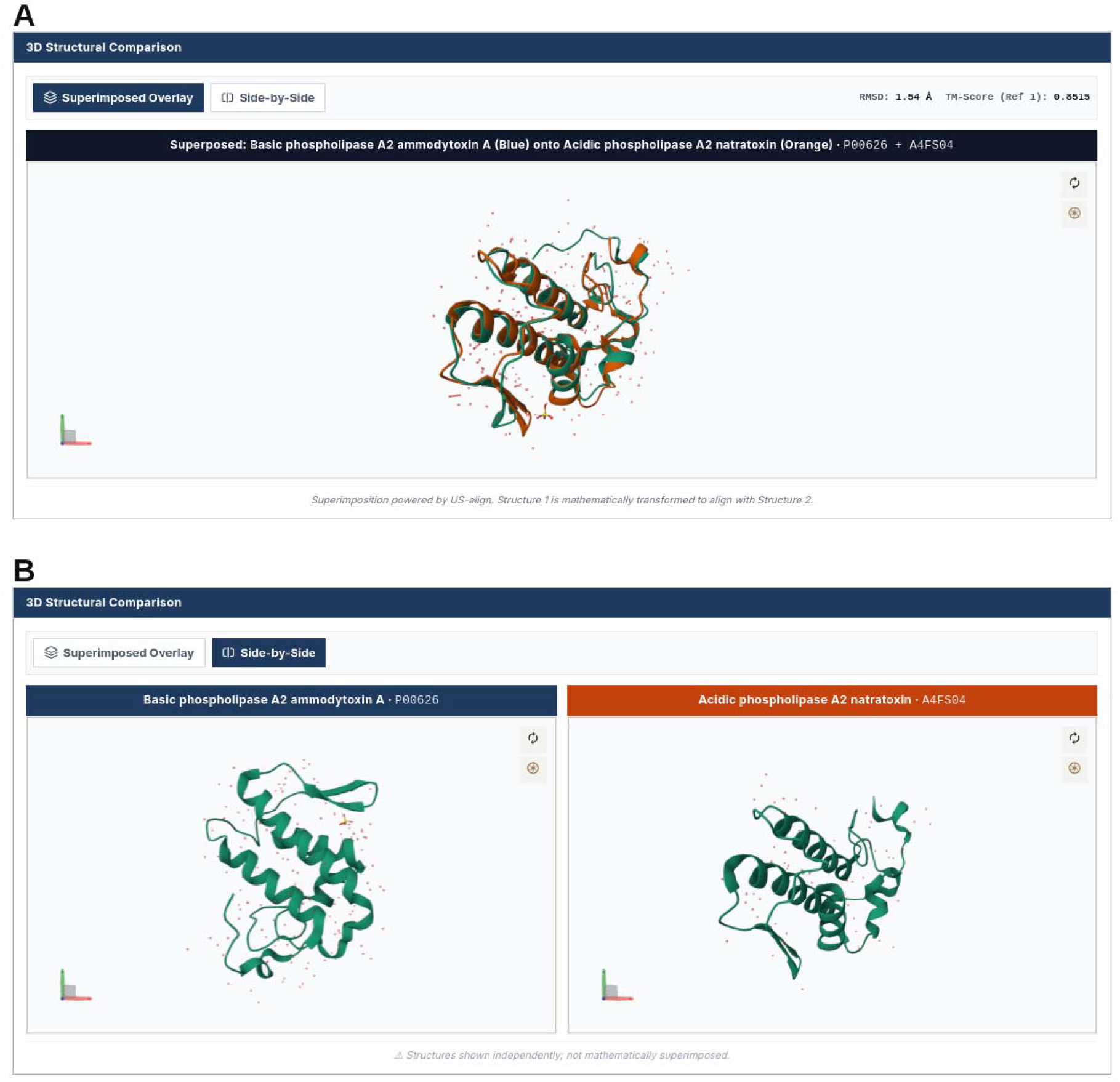
3D structural comparison. (A) Superimposition (RMSD 1.54 Å, TM-score 0.8515). (B) Side-by-side view.

The user can access the guide provided in the index section for clarity on the usage of the database and tools provided.

## 4. ILLUSTRATIVE USE CASES

ToxINsight combines a distribution atlas, a searchable toxin database with taxonomy browsing, a personal workspace, and tools for sequence alignment, pairwise comparison and export. These modules can be combined in different ways depending on the specific research question. We describe six illustrative use cases, covering structure-based design, comparison of predicted and experimental models, regional antivenom design, comparison of toxin variants, dataset preparation for machine learning, and the toxin repertoire of a single species. Figure 10 shows the path each use case takes through the platform modules, and the figures that follow show the corresponding screens.

1. **Prioritizing structures as scaffolds for structure-based drug design. A** user browses the Atlas for a snake venom family/species/region of interest, retrieves structures for all matching entries individually, and uses the pairwise comparison tool to identify structurally distinct scaffolds sharing the same reported target. They can now be taken as candidates for structure-based design or as templates for peptide or drug engineering for the region of interest.
2. **Providing missing or comparative structures with ESMFold-derived models.** Even if the structural data is present for the toxin entry, models predicted from ESMFold can still provide useful insights into structural features, as ESMFold works by predicting protein structures from sequence information using a large protein language model (ESM-2) instead of relying on evolutionary information from homologous sequences. This is a major difference from AlphaFold3-predicted models, which depend on multiple sequence alignments (MSAs) and homologous models to capture co-evolutionary signals. Therefore, ESMFold needs no MSA and can model sequences with few to no homologs. The difference in their algorithms means that both AlphaFold3 and ESMFold provide slightly different structures, which could be used for comparative analysis and feature recognition.
3. **Cross-species conserved region mapping for regional antivenom design.** Antivenoms are raised against venoms of regionally important snakes, and their effectiveness varies between regions because venom composition varies between populations and species. A user can therefore start from a region of interest, or directly from a toxin family across several genera or species, and use ToxINsight to locate conserved regions that can be evaluated as candidate cross-reactive epitopes for broad-spectrum antivenom development. Starting from the Distribution Atlas, a user selects a country or region and sees which venomous species and families are reported there. For example, selecting India lists 24 venomous species (15 Elapidae, 8 Viperidae and 1 Homalopsidae), including the “big four” medically important species (*Naja naja*, *Bungarus caeruleus*, *Daboia russelii* and *Echis carinatus*) (Figure 11A). The user can retrieve the toxin entries for these species, save them to a workspace collection (Figure 11B) and run a multiple sequence alignment within a toxin family (Figure 11C). We illustrate this with phospholipase A2 (PLA2), a family present in both elapid and viperid venoms. The alignment of one PLA2 entry from each of the four species shows that all share the calcium-binding loop (YGCYCG) and the catalytic site region (CC-x-x-HD-x-CY), whereas the sequences diverge in other regions. Regions that are conserved across the local species are candidates for cross-reactive epitopes or inhibitor targets, and divergent regions indicate where an existing antivenom may be less likely to neutralise a local venom. Sequence conservation alone does not show that a region is accessible or immunogenic, so candidate regions should be examined on the 3D structures and tested experimentally. Sequences (FASTA) can be exported in bulk, and structures can be downloaded from each entry page, for downstream epitope analysis. The example uses one entry per species and reflects reported entries, not the abundance of each toxin in the venom.
4. **Relating differences between toxin variants to functional sites.** Given that two orthologous or paralogous toxins can be saved individually in the workspace (Figure 12A), a user applies the pairwise comparison tool to identify where they differ in sequence, physicochemical properties and structure, and relates these differences to known functional residues. For example, α-cobratoxin from *Naja kaouthia* (P01391), the target of the peptide neutralisers^50^, and Long neurotoxin 3 from *Naja naja* (P25671) are both 71 residues long and 97.2% identical, differing only by two Asn→Asp substitutions at positions 61 and 63 (Figure 12C). These lower the predicted isoelectric point from 8.60 to 8.11 and the net charge from 3.75 to 1.75 (Figure 12B). Neither substitution is among the toxin residues in contact with the peptide inhibitor in the reported crystal structure (D27, F29, R33, R36 and F65, plus the P7 backbone), which suggests that such peptides might also recognise the Indian cobra toxin.
5. **Bulk download for machine-learning approaches.** A researcher building predictive ML models for snake venom toxins can select an entire family, a taxonomic group or a custom set of entries and export sequences (FASTA) and tabular metadata (CSV) in a single download. Three-dimensional structures (PDB) are available for each entry from its toxin page. Each record carries the toxin family, source species, length and computed physicochemical properties, so the data can be turned into labelled datasets for tasks such as toxin family classification, property prediction, or protein language model embeddings and fine-tuning. Because each entry records whether it is reviewed (Swiss-Prot) or unreviewed, and whether its structure is experimental or predicted, users can filter to higher-confidence subsets before training. Users should cluster sequences by identity before splitting into training and test sets, as closely related toxins can otherwise leak between them, and should account for the uneven family sizes in the database. (Figure 13)
6. **Species toxin repertoire.** A researcher interested in the venom of a particular snake, for example, the king cobra (*Ophiophagus hannah*), can search ToxINsight by scientific or common name, or open the species through the taxonomy browser, and retrieve all toxin entries reported for it. Filtering by toxin family shows which families are represented, such as three-finger toxins, phospholipases A2 or metalloproteinases. The entries can be saved to a workspace collection, where accession, name, species, family and length are listed together, and exported as CSV to tabulate the number of entries per family. Within a collection, entries of the same family can be sent to multiple sequence alignment to see how related variants differ. Any two entries can be compared on physicochemical properties, structure and global sequence alignment. A downloadable PDF report for each entry gives its identification, functional description, physicochemical properties, available 3D structures, Gene Ontology annotations and references. Together, these steps let researchers assemble and analyse the reported toxin repertoire of one species in one place. The repertoire reflects the entries reported in the database, not the relative abundance of each toxin in the venom.

**Figure 10.**
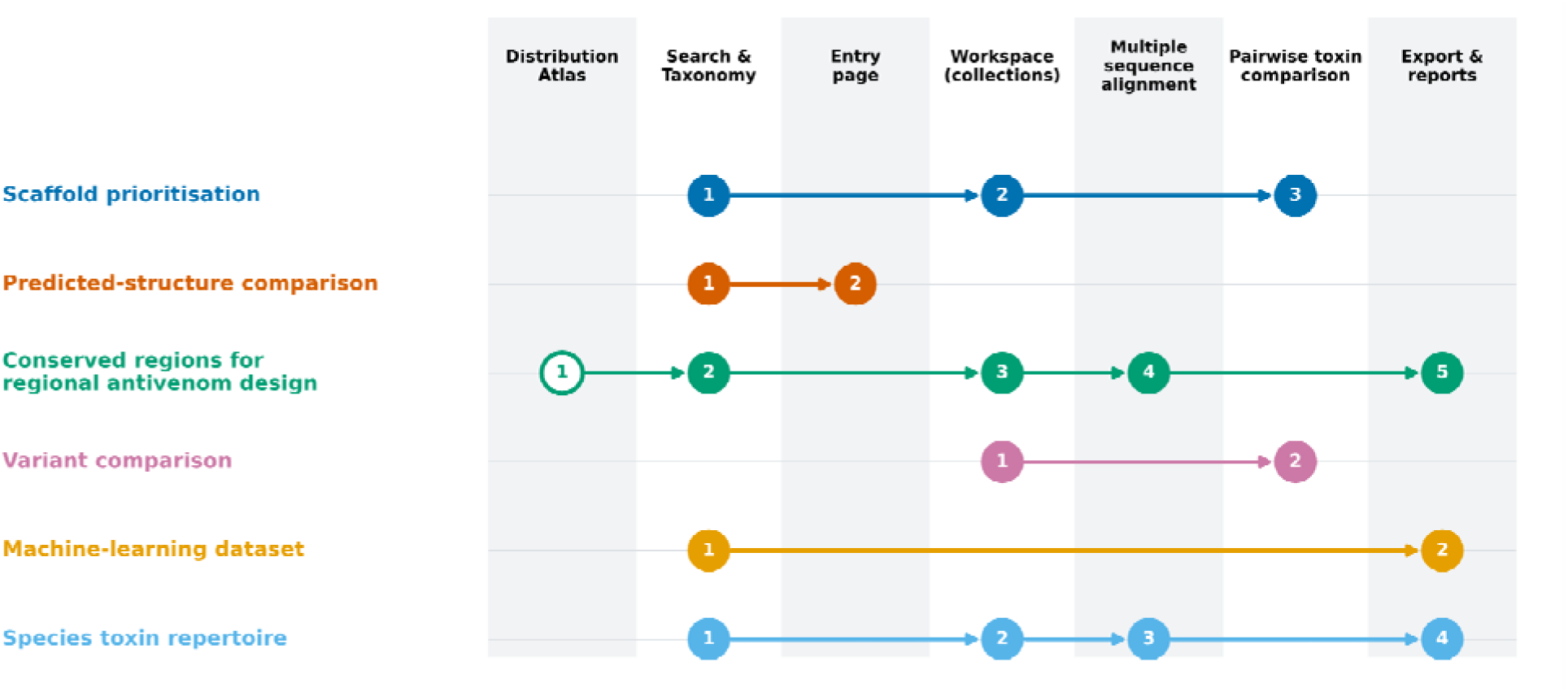
ToxINsight supports six research workflows through a small set of connected modules. Rows, from top to bottom, correspond to the use cases described in this section: (1) scaffold prioritisation for structure-based design, (2) predicted-structure comparison, (3) cross-species conserved-region mapping for regional antivenom design, (4) comparison of toxin variants, (5) machine-learning dataset preparation, and (6) species toxin repertoire. Columns are the platform modules: Distribution Atlas, Search and Taxonomy, Entry page, Workspace, Multiple Sequence Alignment, Pairwise Toxin Comparison, and Export and Reports. The numbered circles give the order in which a user moves through the modules for each use case, and the arrows show the direction of the workflow. The open circle in use case 3 marks the Atlas step, which is optional when starting from a toxin family rather than a region.

**Figure 11.**
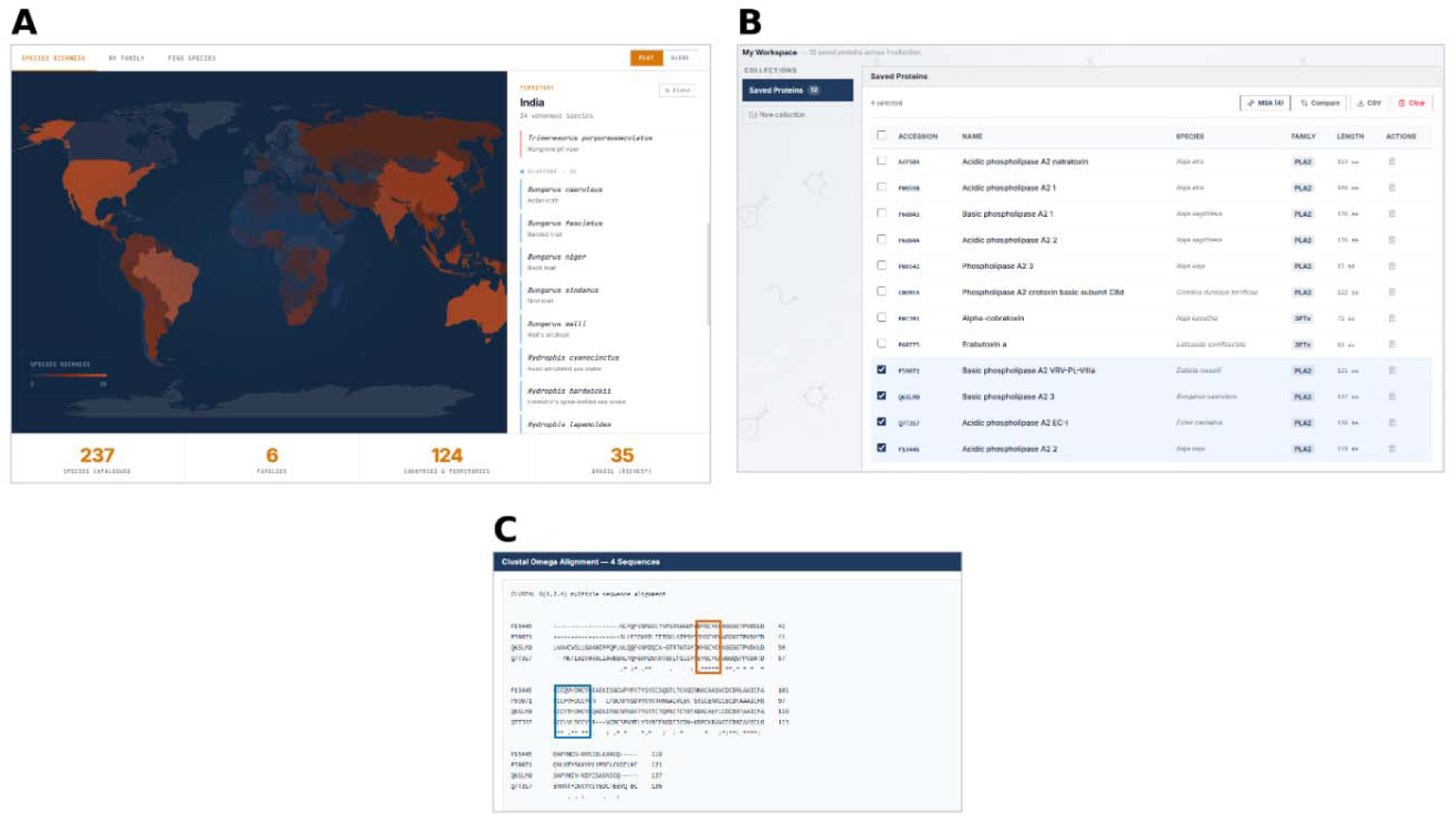
Mapping conserved regions in venom toxins for regional antivenom design. (A) Selecting India in the Distribution Atlas lists its 24 venomous species by family in the side panel; scrolling would show the Viperidae, Elapidae and Homalopsidae. (B) Phospholipase A2 entries for Naja naja (P15445), Daboia russelii (P59071), Bungarus caeruleus (Q6SLM0) and Echis carinatus (Q7T3S7) are saved to a Workspace collection and sent to alignment. (C) Clustal Omega multiple sequence alignment of the four sequences, shown as stored in the database. Boxes mark the calcium-binding loop (orange) and the active-site region (blue), both highly conserved across the four species.

**Figure 12.**
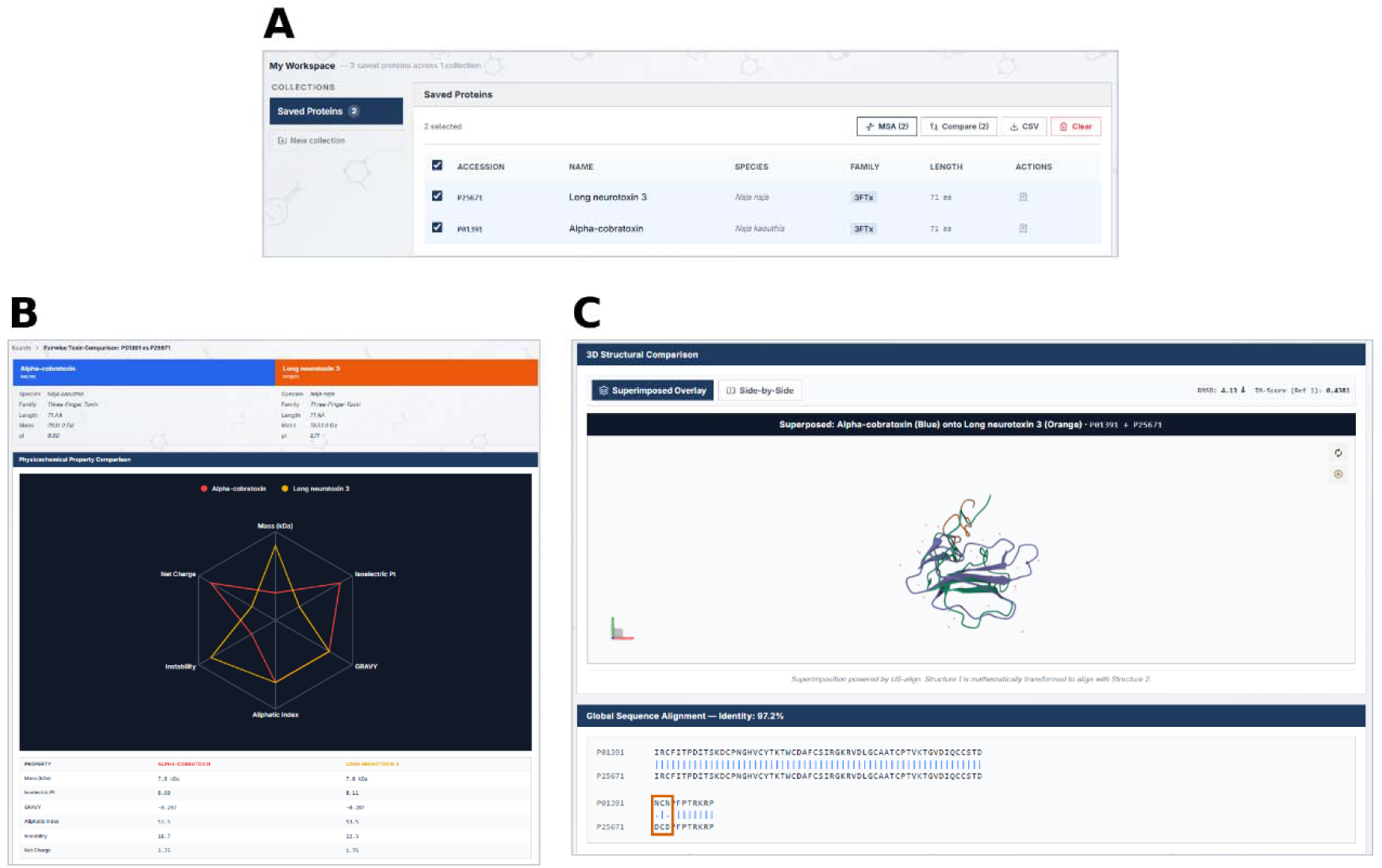
Comparing two close long-chain α-neurotoxin variants with the Pairwise Toxin Comparison tool. (A) α-Cobratoxin from *Naja kaouthia* (P01391) and Long neurotoxin 3 from *Naja naja* (P25671) are saved to a Workspace collection and sent for comparison. (B) Physicochemical comparison: both toxins are 71 residues long with identical GRAVY and aliphatic index, and differ in isoelectric point (8.60 and 8.11), net charge (3.75 and 1.75), and instability index (18.7 and 22.3). Radar axes are scaled between the two toxins. (C) Structural superposition and global sequence alignment. The sequences are 97.2% identical and differ at two positions, both Asn→Asp (boxed, residues 61 and 63).

**Figure 13.**
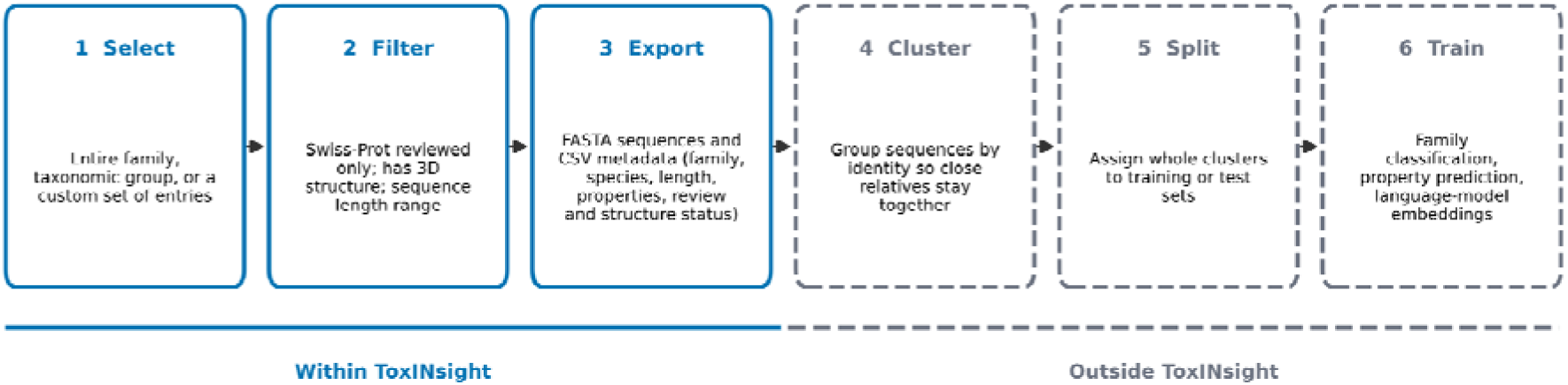
Building a labelled dataset for machine learning from ToxINsight. Steps 1 to 3 are carried out in ToxINsight and steps 4 to 6 in the user’s own pipeline. (1) Select: the user chooses an entire toxin family, a taxonomic group or a custom set of entries. (2) Filter: entries are restricted to higher-confidence subsets, using Swiss-Prot reviewed status, availability of a 3D structure and a sequence length range. (3) Export: sequences (FASTA) and tabular metadata (CSV) are downloaded in bulk. The CSV records toxin family, source species, length, computed physicochemical properties, review status and structure type. (4) Cluster: sequences are grouped by identity so that closely related toxins stay together. (5) Split: whole clusters are assigned to training or test sets, which prevents leakage between them. (6) Train: models are trained or fine-tuned for tasks such as toxin family classification, property prediction, or protein language model embeddings.

## 5. CONCLUSIONS AND FUTURE DIRECTIONS

ToxINsight brings together curated sequence, structural coverage, reported ligand/target annotation, and a built-in comparative toolset (MSA and pairwise structural/physicochemical comparison) for snake venom toxins in a single, freely accessible resource. By combining experimental structures, AlphaFold DB predictions, and ESMFold-based gap-filling, ToxINsight ensures that every toxin entry has an associated three-dimensional model available for immediate structural analysis, removing a common limitation in snake venom research. We anticipate that ToxINsight will be useful across structure–activity relationship analysis, comparative venomics, antivenom design, and venom-derived peptide drug discovery.

## Author Contributions

B.S. ideated and supervised the project. A.G. and T.V.L.K. curated the toxin data. T.V.L.K. built the databases and implemented the analysis tools. A.G. wrote the manuscript. All authors reviewed and approved the final manuscript.

## Notes

The authors declare no competing financial interests.

## Abbreviations

3FTx: three-finger toxin
PLA2: phospholipase A2
SVMP: snake venom metalloproteinase
SVSP: snake venom serine protease
CTL: C-type lectin/lectin-like protein
KUN: Kunitz-type protease inhibitor
DIS: disintegrin
CRISP: cysteine-rich secretory protein
LAAO: L-amino-acid oxidase
NP: natriuretic peptide
PDB: Protein Data Bank
MSA: multiple sequence alignment
RMSD: root-mean-square deviation.

